# Molecular determinants of WNT9b responsiveness in nephron progenitor cells

**DOI:** 10.1101/328765

**Authors:** Kyle K. Dickinson, Leah C. Hammond, Courtney M. Karner, Nicholas D. Hastie, Thomas J. Carroll, Paul R. Goodyer

## Abstract

Primed nephron progenitor cells (NPCs) appear in metanephric mesenchyme by Ell.5 and differentiate in response to the inductive WNT9b signal from the ureteric bud. However, the NPC WNT-receptor complex is unknown. We obtained M15 cells from E10.5 mesonephric mesenchyme and systematically analyzed components required for canonical WNT9b-responsiveness. When M15 cells were transfected with a (β-catenin luciferase reporter plasmid, exposure to recombinant WNT9b resulted in minimal luciferase activity. We then analyzed mRNA-expression of WNT-pathway components and identified *Fzd1-6* and *Lrp6* transcripts but not *RSPO1.* When M15 cells were treated with recombinant RSPO1 the response to transfected WNT9b was augmented 4.8-fold. Co-transfection of M15 cells with *Fzd5* (but no other *Fzd* family member) further increased the WNT9b signal to 16.8-fold and siRNA knockdown of *Fzd5* reduced the signal by 52%. Knockdown of *Lrp6* resulted in 60% WNT9b signal reduction. We confirmed *Fzd5, Lrp6* and *RSPO1* rrtRNA expression in CITED1(+) NPCs from E15.5 embryonic mouse kidney. Thus, while many WNT signaling-pathway components are present by E10.5, optimum responsiveness of Ell.5 cap mesenchyme requires that NPCs acquire RSPO1, FZD5 and LRP6.

**Summary Statement:** Responsiveness to the inductive WMT9b signal from ureteric bud is crucial for nephrogenesis. Here we analyze the molecules needed to prime nephron progenitor cells in embryonic mouse kidney.

## Introduction

The mammalian kidneys are derived from progenitor cells in the embryonic intermediate mesoderm, expressing the transcription factor, OSR1. Murine fate mapping studies at embryonic day E7.5 indicate that cells labeled by the *Osr1* promoter at E7.5 give rise to all elements of the murine metanephric kidney (Mugford et al., 2008) and *Osr1* knockout mice are anephric (James et al., 2006; Lan et al, 2011). By about E8.5-E9, the OSR1(+) kidney progenitor cells reach a critical fork in the road. Some of the cells are transformed into PAX2(+)/GATA3(+) polarized epithelia, forming paired nephric ducts that elongate caudally down the embryo (Bouchard et al., 2002). A second subset of cells retain the mesenchymal phenotype and begin to express the Wilms’ tumor gene, WT1 (Narlis et al., 2007; Wilm and Munoz-Chapuli, 2016). The columns of WT1(+) cells flanking each nephric duct are committed to the nephron progenitor fate; *Wt1* knockout mice are completely anephric (Schedl and Hastie, 1998). Development of the metanephric kidney begins in earnest when ureteric buds emerge from each nephric duct (E10.5), begins to arborize as it grows into the adjacent column of metanephric mesenchyme and induces local nephron progenitors to begin nephrogenesis.

In the 1950s, Grobstein showed that fragments of metanephric mesenchyme can generate renal tubular structures if they are co-cultured with certain inductive tissues such as embryonic spinal cord that can substitute for the ureteric bud (Grobstein, 1956). This seminal discovery showed that the metanephric mesenchyme contains a population of committed nephron progenitor cells that can unleash the differentiation cascade of nephrogenesis if exposed to the proper signals from ureteric bud. Key observations by Herzlinger (Herzlinger et al., 1994) and Carroll (Carroll et al., 2005; Karner et al., 2011) established the canonical WNT9b/β-catenin signaling pathway as the central mechanism by which the ureteric bud initiates nephrogenesis. WNT9b secreted by the ureteric bud is essential for the early inductive events; mice bearing a (3-catenin-responsive reporter transgene exhibit intense canonical WNT signaling activity in the layer of progenitor cells immediately adjacent to each ureteric bud branch tip (Schmidt-Ott and Barasch, 2008; Iglesias et al., 2014).

It is unclear when nephron progenitors become competent to respond to the inductive WNT signal, but WT1 is crucial. In humans, biallelic mutations of the *WT1* gene lead to a developmentally arrested clone of cells (“nephrogenic rests”) which lack canonical WNT/(3-catenin activity (unlike their heterozygous sister cells) and are apparently unable to respond to inductive signals from the ureteric bud (Fukuzawa et al., 2009). We discovered that this is accomplished by WT1 suppression of EZH2, de-repressing epigenetically silenced genes of the differentiation cascade (Akpa et al., 2015). Within a day of WT1 appearance and prior to arrival of the ureteric bud (E10.5-Ell), maturing NPC express a panel of genes, including retinoic acid receptor-alpha, cadherin 11 and CD24 (Challen et al., 2004; Iglesias et al., 2014). However, the stage at which they are fully competent to respond to the WNT9b signal is unknown. Furthermore, the molecular basis for WNT9b responsiveness in nephron progenitor cells is unknown.

The canonical WNT signaling pathway is full of redundancies. Here we take a systematic approach to identifying the crucial components of the WNT9b signaling pathway in embryonic mouse kidney.

## Results

### M15 Cells

From fate mapping studies in embryonic mouse kidney, a committed lineage of nephron progenitor cells (NPC) emerges from the OSR1(+) intermediate mesoderm as early as embryonic day E7.5 (Mugford et al., 2008). These cells express the Wilms Tumour transcription factor, WT1. To model early events that render NPCs responsive to the inductive WNT9b signal from ureteric bud, we analyzed the M15 cell line, derived from E10.5 mesonephric mesenchyme of mice bearing the large T polyoma virus protein under the control of an early viral enhancer (Larsson et al., 1995). These cells are thought to represent the NPC phenotype one day prior to arrival of the ureteric bud at Ell.5. We confirmed expression of WT1 in M15 cells by RT-PCR (Fig 1a) and Western immunoblotting **(Fig 1b).** We then screened the M15 cells for mRNA expression (RT-PCR) of candidate genes in the canonical WNT-βcatenin signaling pathway (Table 1). We identified expression of β-Catenin, *Lrp6, Lgr4/6* and *Fzd1-6.* Notably absent were *RSPO1* and 3, *Fzd7-10, Lrp5* and *Lgr5.*

**Fig 1A.**
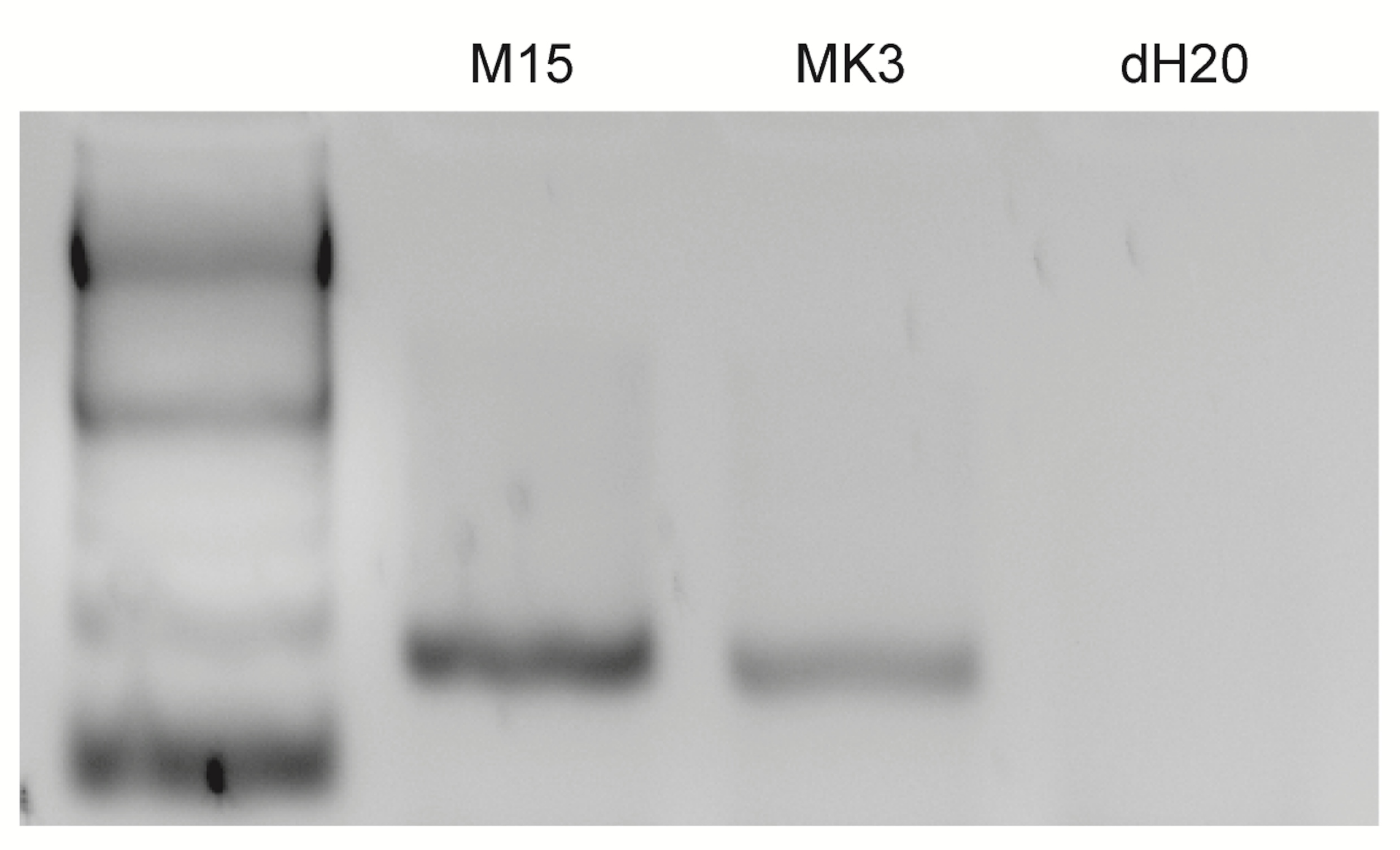
WT1 is expressed in M15 cells. mRNA from E10.5 mouse metanephric mesenchyme (M15 cells) was analyzed by RT-PCR for Wt1 mRNA expression in M15 cells and MK3 (positive control) (Potter) cells vs water blank.

**Fig 1B:**
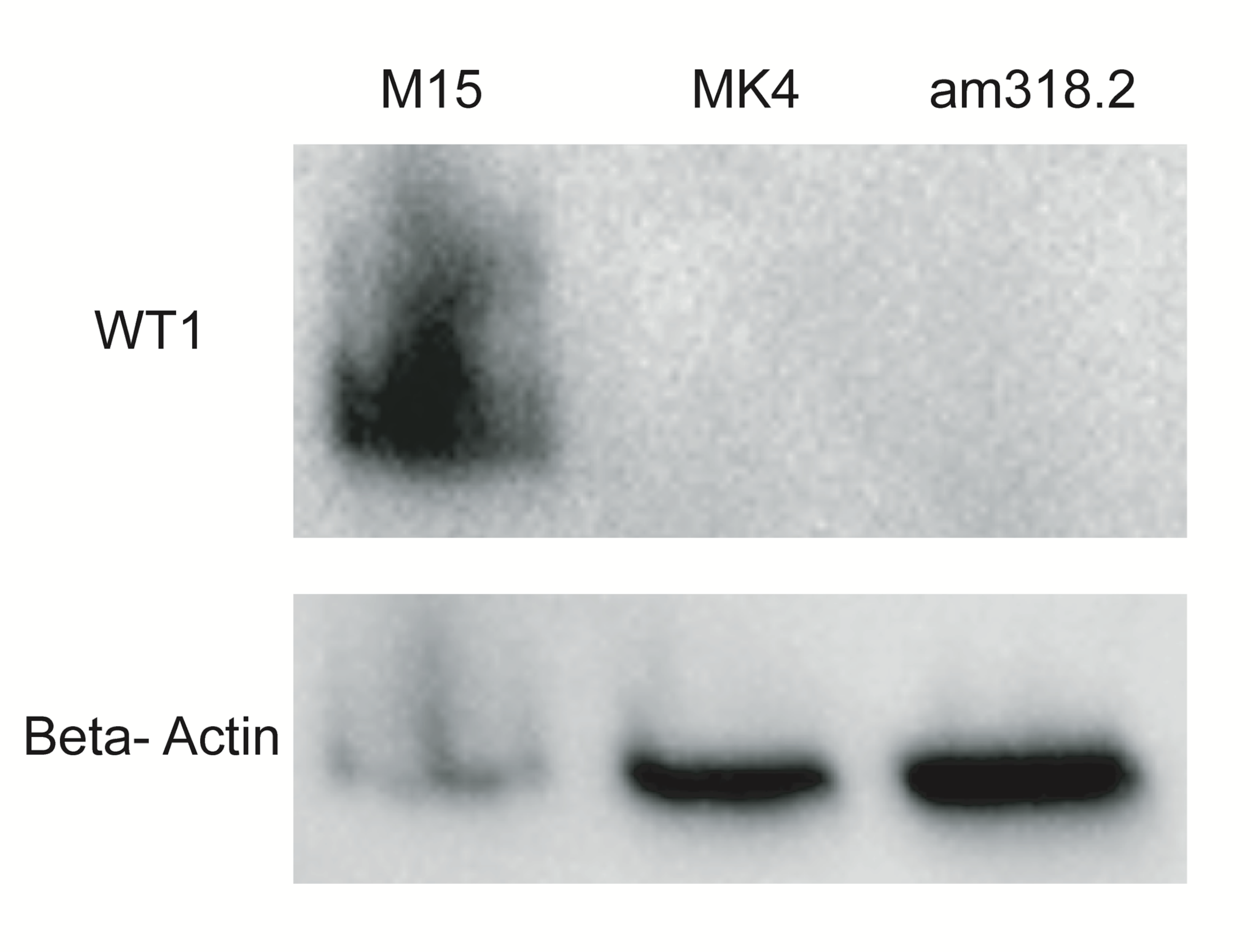
WT1 is expressed in M15 cells. Lysates of M15 cells vs E 14.5 MK4 (negative control) (Potter) or am318.2 mesenchymal stem cells from 20-week gestation human amniotic fluid (Murielle) were analyzed by Western immunoblotting for WT1 protein (upper panel) and Beta actin (lower panel).

**Table I.**
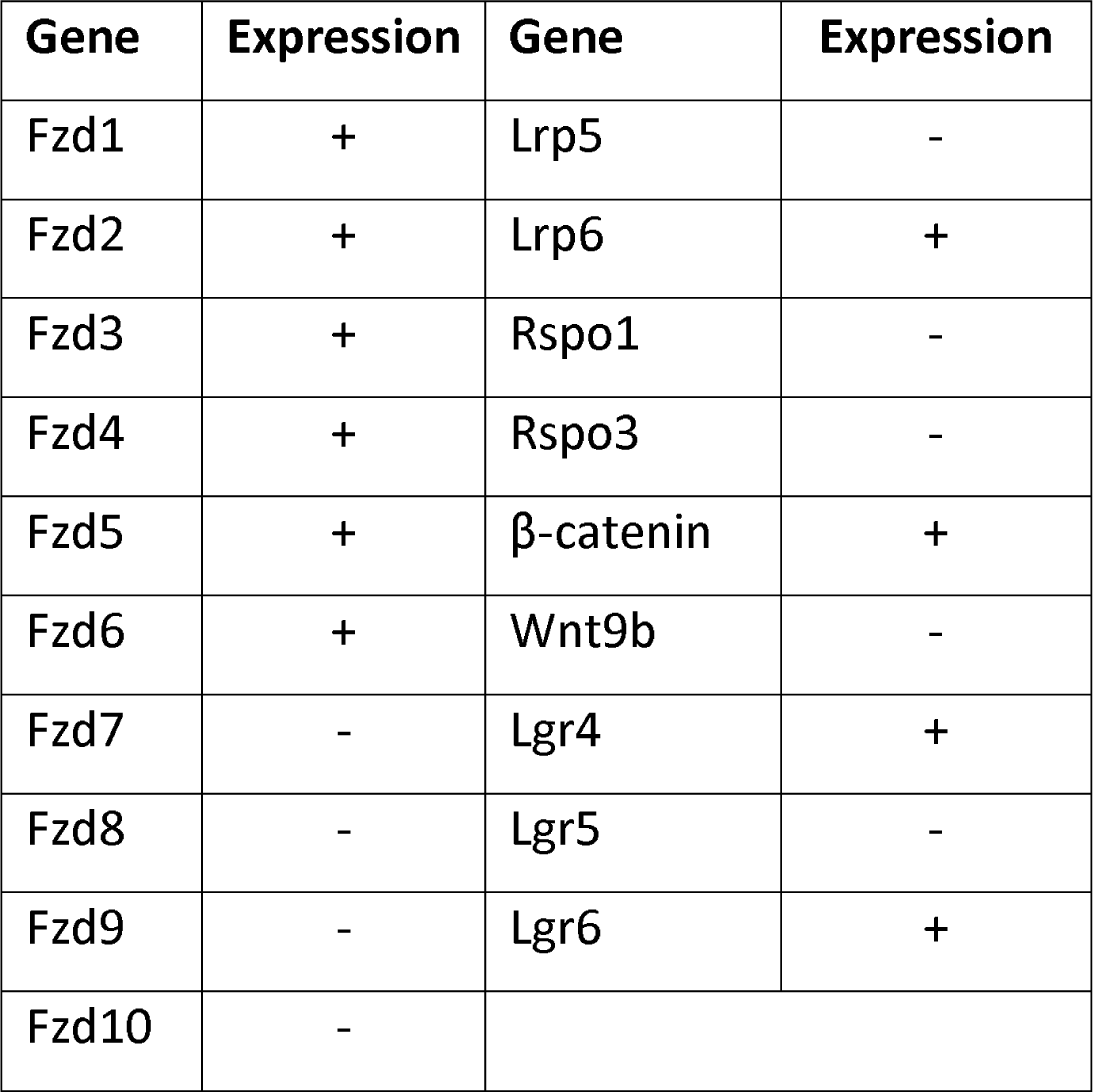
mRNA expression of WNT/P-catenin pathway components in M15 cells.

### M15 cells are unresponsive to external WNT9b

To ascertain whether M15 cells are primed to respond to a WNT9b signal, we transiently transfected the cells with a (3-Catenin/luciferase reporter (TOPFlash) and exposed them to recombinant WNT9B protein at concentrations from 50-400 ng/ml but detected only minimal response (1.05-fold) **(Fig 2).**

**Fig 2:**
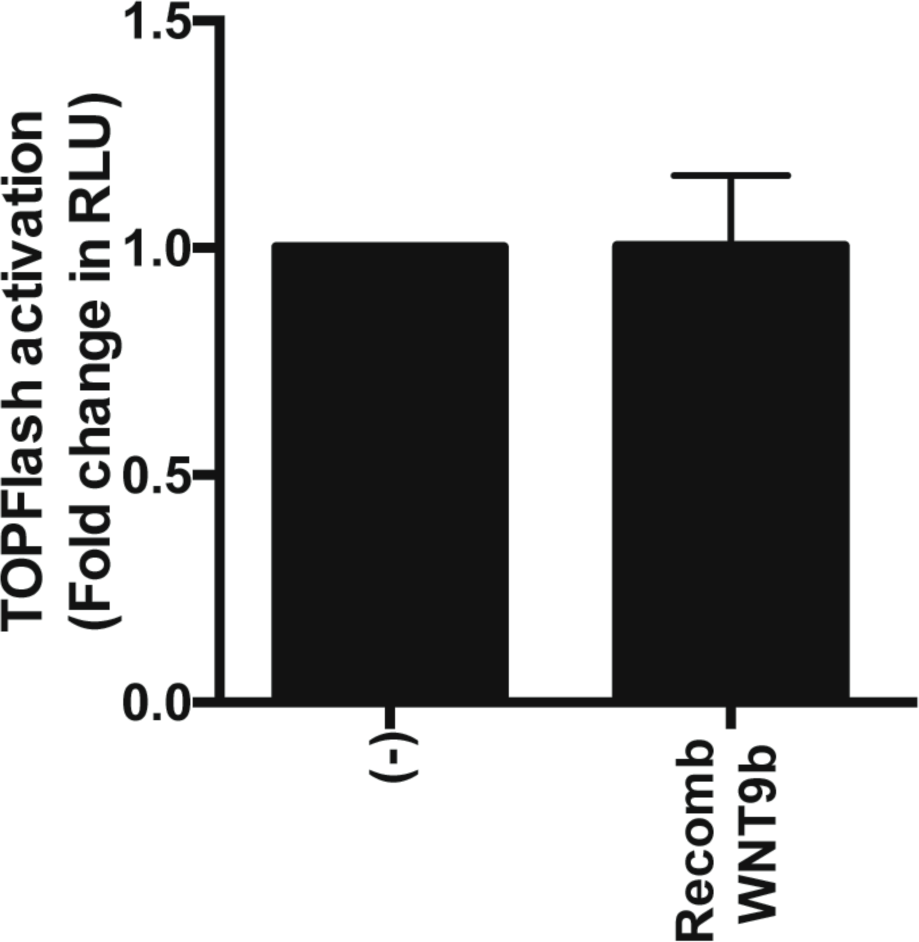
WNT-responsiveness of M15 cells. M15 cells were transiently transfected with (3-catenin-luciferase reporter (8X SuperTOPFlash) and Renilla-luciferase reporter. The cells were exposed to recombinant WNT9b (50 ng/ml). After 48 hours, TOPFlash to Renilla signal (RLU) was measured in a luminometer. An unpaired two-tailed Welch’s t-test was performed, (ns) p=0.98.

### RSPO1 enhances responsiveness of M15 cells to WNT9b

Since, M15 cells lack both R-spondins known to be expressed in nephron progenitor cells of embryonic mouse kidney cap mesenchyme (GUDMAP), we reasoned that M15 cell WNT responsiveness might be limited by stability of the WNT receptor complex at the cell surface (Binnerts et al., 2007; McMahon et al., 2008; Harding et al., 2011). To test this hypothesis, we first transfected M15 cells with (β-Catenin/luciferase reporter (TOPFlash) as above and assessed the response to a co-transfected WNT9b expression plasmid. As seen in Fig 3, a significant (5-fold) increase in luciferase activity was seen. We then added recombinant RSPO1 (200 ng/ml) or R-SP03 (200 ng/ml). Addition of either R-spondin further increased the signal to 22- and 27-fold above baseline, respectively (p<0.0001) (Fig 3). Preliminary dose-response studies showed that no further signal increase was obtained with higher concentrations of either R-spondin. To dissect the importance of other canonical WNT pathway components, we added *Wnt9b* plasmid and recombinant RSPO1 (200 ng/ml) in all subsequent experiments.

**Fig 3:**
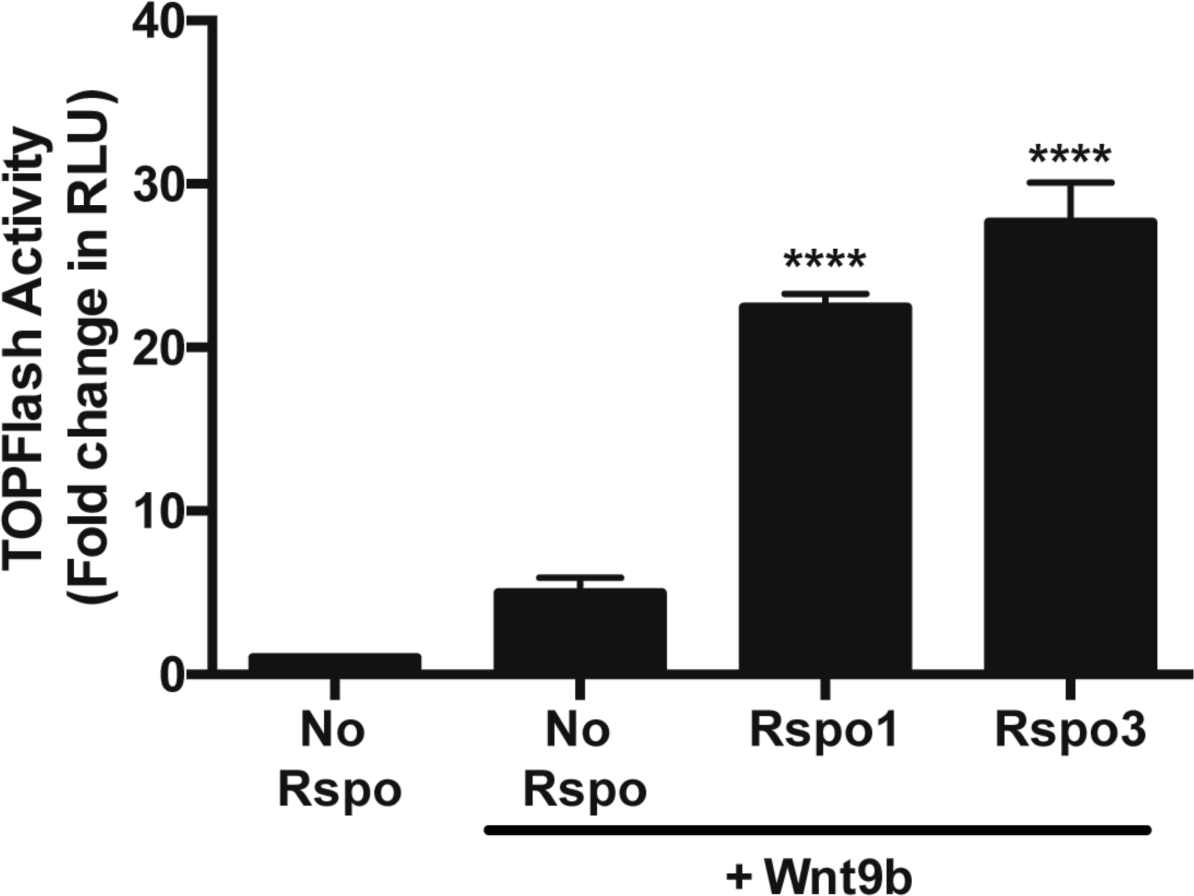
Effect of recombinant RSP01 on responsiveness of M15 cells to WNT9b. M15 cells were transfected with 8X SuperTOPFlash, Renilla and Wnt9b plasmids and cultured for 24 hours; recombinant RSP01 or RSP03 (200 ng/ml) were added for an additional 24 hours and TOPFlash to Renilla signal was measured. A one-way ANOVA followed by a Dunnett correction for multiple comparisons was performed. (****) = p <0.0001.

### Frizzled receptors in cap mesenchyme

The Fzd receptor mediating the inductive WNT9b signal during nephrogenesis is unknown. To identify candidate Frizzled receptors transducing the WNT9b response in nephron progenitors, we performed in situ hybridization for the various Frizzled family members (*Fzd1-10*) in Ell.5 mouse kidney, except *Fzd9* which was unsuccessful. As seen in Fig 4, several Frizzled family members (*Fzd1, Fzd2, Fzd3, Fzd5 and Fzd7)* are strongly expressed in the cap mesenchyme (but not in the ureteric bud); this pattern contrasts with *Fzds* with weak non-specific expression patterns (*Fzd 4* and *FzdlO*) or strong expression restricted to ureteric bud branch tips (*Fzd6* and *Fzd8).*

**Fig 4:**
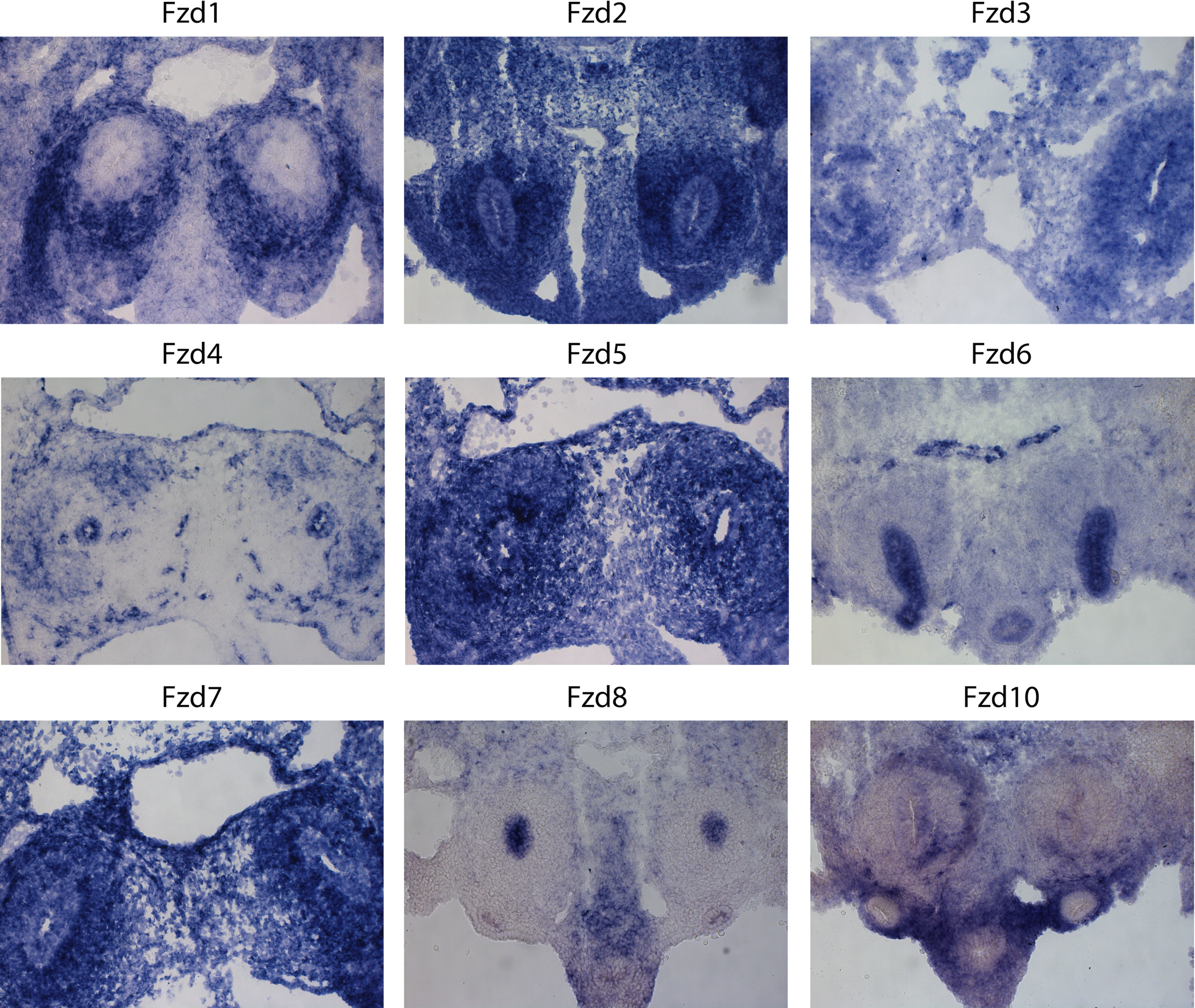
Frizzled (FZD) mRNA expression in embryonic day E11.5 mouse kidney. Cryosections of E11.5 mouse kidneys were assessed by in situ hybridization using riboprobes for Fzd 1-10, except Fzd9 which was unsuccessful for technical reasons.

### Transfection of M15 cells with *Fzd5* enhances WNT9b responsiveness

To ascertain whether one of the FZD receptors is rate limiting in M15 cells, we separately transfected each member of the *Fzd* receptors family (*Fzds 1-10*) into M15 cells expressing the (β-Catenin/luciferase reporter (TOPFlash). All cells were co-transfected with Wnt9b plasmid and exposed to recombinant RSPO1 (200 ng/ml) in each experiment. As seen in Fig 5A, the only Fzd which significantly augmented WNT9b-induced TOPFlash signal was Fzd5. When M15 cells were cotransfected with *Fzd5,* activity of the canonical WNT/β-Catenin reporter was increased 3.5-fold (p=0.0002). We then performed similar experiments in M15 cells co-transfected with an siRNA targeting Fzd5, previously shown to knock down Fzd5 expression level by 70%. As seen in Fig 5B, presence of the *Fzd5* siRNA reduced WNT9b-dependent TOPFlash activity by 52% (p=0.005).

**Fig 5A:**
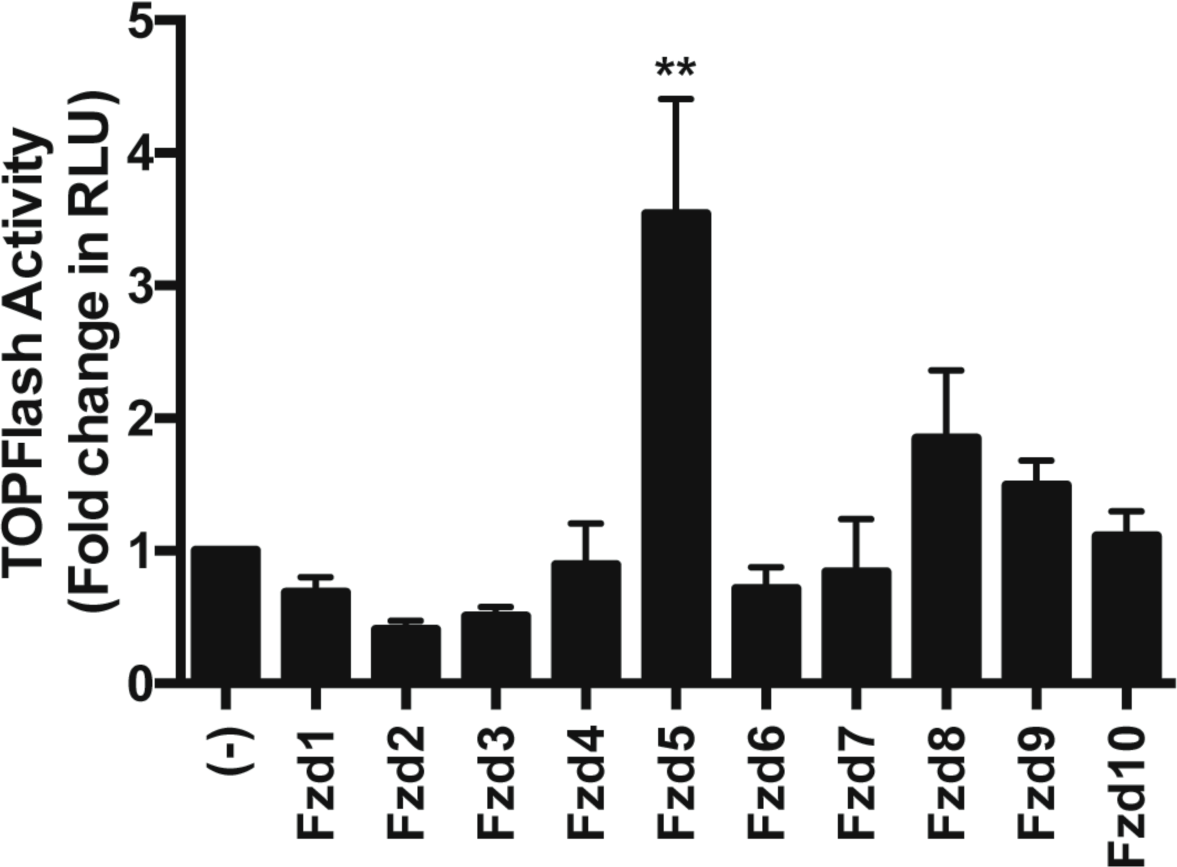
Effect of FZD expression on WNT9b responsiveness in M15 cells. M15 cells were transiently transfected with [3-catenin-luciferase reporter (8X SuperTOPFlash), Renilla-luciferase reporter, Wnt9b-expression vector and various Fzd 1-10 expression plasmids in the presence of recombinant RSP01 (200 ng/ml). TOPFlash to Renilla signal was measured after 48 hours. A one-way ANOVA followed by a Dunnett correction for multiple comparisons was performed. (**) p=0.0002.

**Fig 5B:**
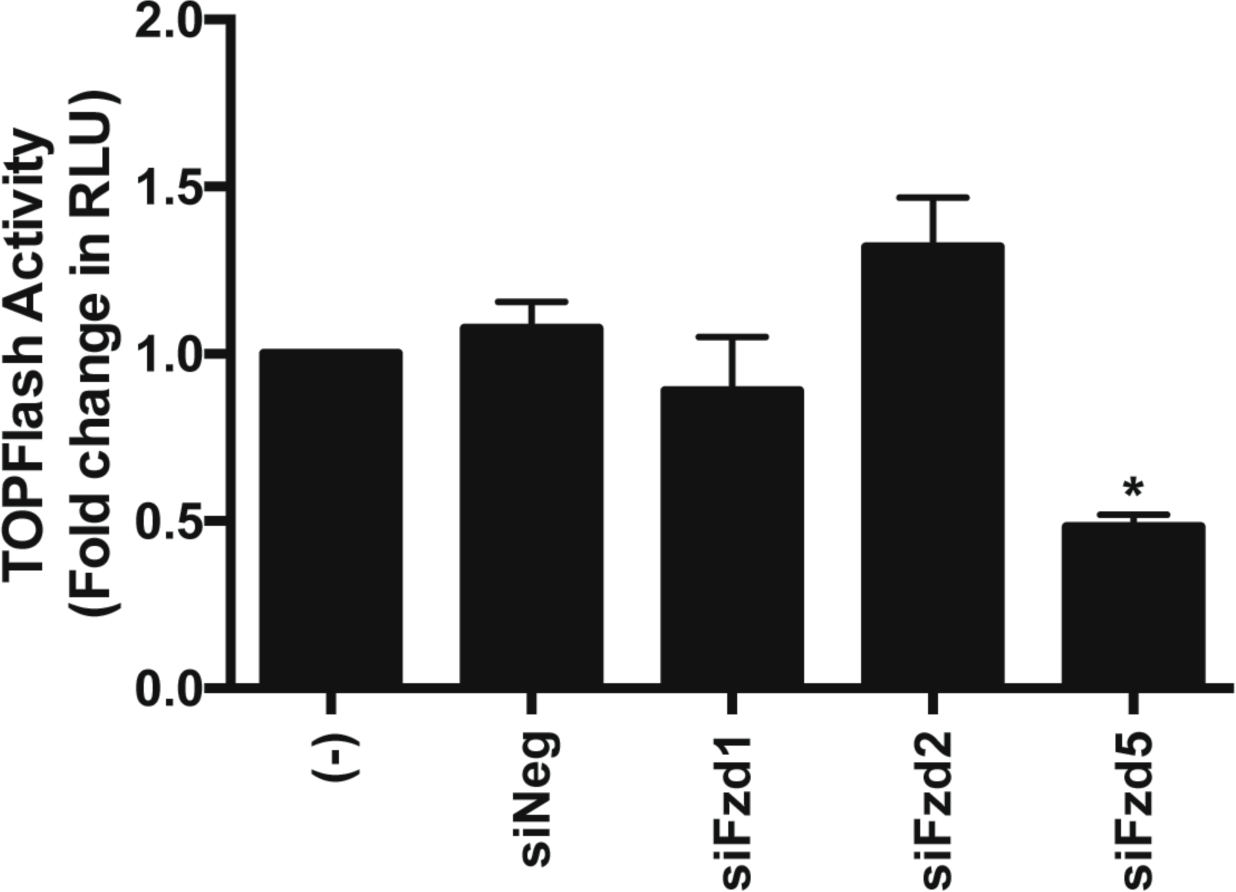
Effect of FZD expression on WNT9b responsiveness in M15 cells. M15 cells were transiently transfected with (3-cate nin-luc if erase reporter (8X SuperTOPFlash), Renilla-luciferase reporter, Wnt9b-expression vector and siRNAs targeting Fzdl, Fzd2 or Fzd5 vs a scrambled negative control siRNA in the presence of recombinant RSP01 (200 ng/ml). TOPFlash to Renilla signal was measured. A one-way ANOVA followed by a Dunnett correction for multiple comparisons was performed. (*) p=0.005.

### LRP6 is required for optimal responsiveness of M15 cells to WNT9b

To examine the importance of LRP expression to the canonical WNT-signaling pathway in M15 cells, we transiently transfected M15 cells with WNT9b, β-Catenin/luciferase reporter (TOPFlash) and *Lrp6* siRNA or a scrambled siRNA control. As seen in Fig 6, addition of the LRP6 siRNA reduced WNT9b-dependent TOPFlash signal by 66% (p<0.0001) whereas the scrambled siRNA had no effect. Interestingly, additional co-transfection with LRP5 was unable to rescue WNT9b pathway activity in the presence of LRP6 siRNA. Co-transfection of M15 cells with *Lrp5* (in the absence of siRNA) had no effect on its own.

**Fig 6:**
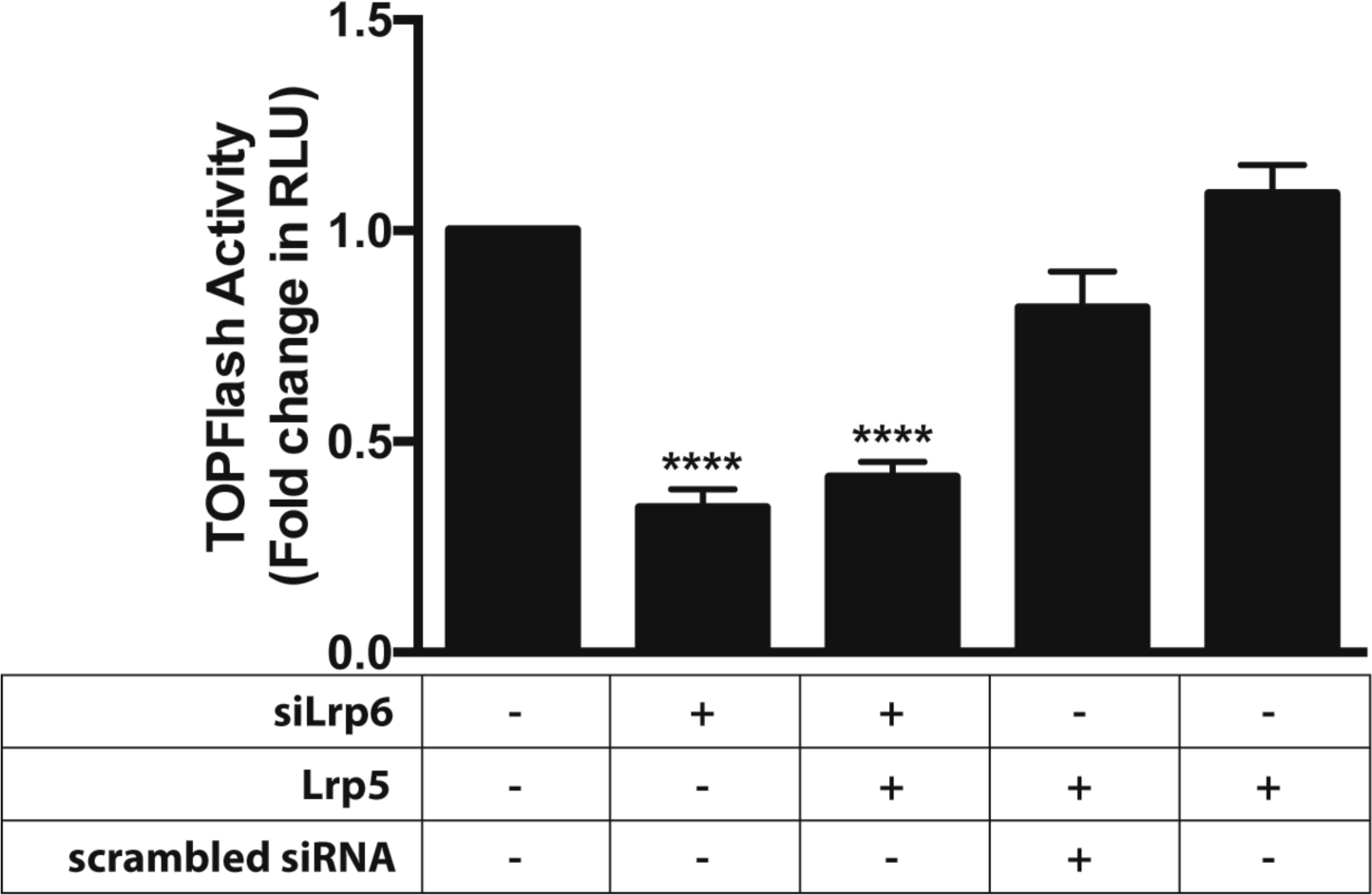
LRP6 is required for optimal responsiveness of M15 cells to WNT9b. M15 cells were transiently transfected with ß-catenin-luciferase reporter (8X SuperTOPFlash), Renilla-luciferase reporter and a Wnt9b expression vector and treated with RSP01 (200 ng/ml). The cells were co-transfected with an siRNA targeting Lrp6 or a scrambled negative control siRNA in the presence of recombinant RSP01 (200 ng/ml). After 48 hours, TOPFlash to Renilla signal was measured. In another experiment, the cells were co-transfected with an Lrp5 expression plasmid to assess its effect on WNT9b pathway activity. A one-way ANOVA followed by a Dunnett correction for multiple comparisons was performed. (****) p<0.0001.

### Responsiveness to extrinsic WNT9b is restored by addition of *Fzd5* and RSPO1

To ascertain whether M15 cell responsiveness to WNT9b could be restored by addition of suboptimal WNT pathway components, we transfected the cells with Fzd5 and treated them with recombinant RSPO1 before assaying TOPFLASI-I/Renilla activity as above. As seen in Fig 7, no response was seen in cells exposed to WNT9b, RSPO1 or *Fzd5* alone. However, the signal was increased 3.3-fold over baseline in M15 cells exposed to recombinant WNT9b and RSPO1. The signal was increased to 11.1-fold over baseline in M15 cells transfected with Fzd5 and then exposed to WNT9b and RSPO1 (p<0.0001) **(Fig 7)**.

**Fig 7:**
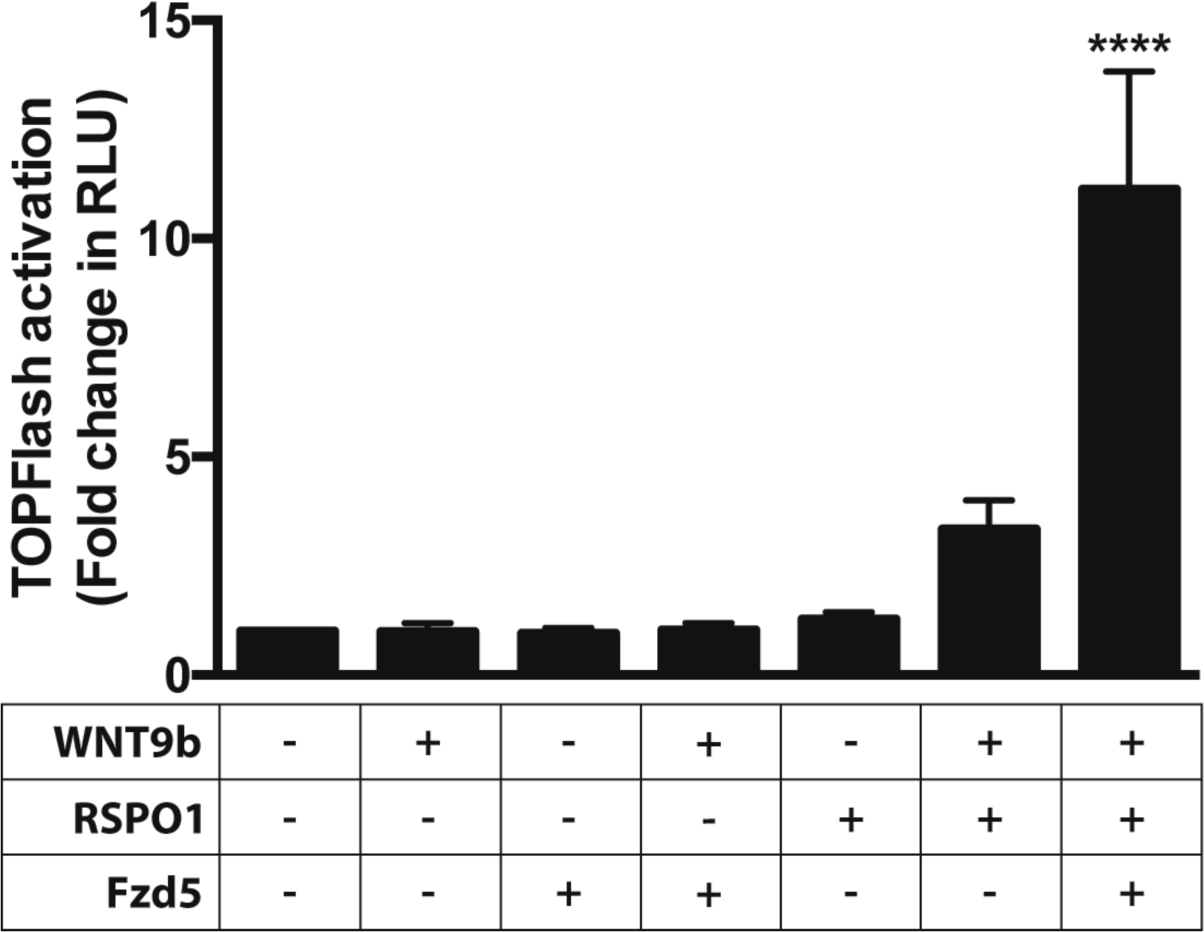
Responsiveness to extrinsic WNT9b is restored by addition of Fzd and RSP01. In all conditions, M15 cells were transiently transfected with (3-catenin-luc if erase reporter (8X SuperTOPFlash) and Renilla-luciferase reporter; in some experiments the cells were co-transfected with Fzd5 expression plasmid. TOPFlash to Renilla signal was measured in the presence or absence of recombinant WNT9b (50 ng/mL) and/or recombinant RSP01 (200 ng/ml). A one-way ANOVA followed by a Dunnett correction for multiple comparisons was performed. (****) = p<0.0001.

### Citedl cells isolated from embryonic mouse kidney express WT1, Fzd5, LRP6 and RSPO1

To identify CITED1 cells in the cap mesenchyme of embryonic mouse kidney, we crossed mice with a floxed TomatoRed transgene to mice bearing a tamoxifen-inducible *Cited1*-driven Cre Recombinase. As seen in Fig 8A, tamoxifen administered to the pregnant mother at E17, activated TomatoRed in progenitor cells of cap mesenchyme **(Fig 8A)** To isolate progenitor cells rapidly after activation of the TomatoRed tag, we digested E15.5 embryonic kidneys from *Citedl*^*Cre*^/TomatoRed mice with collagenase, dispersed the cells into monolayer culture and added tamoxifen (2.5μg/ml) to induce Cre recombinase expression *in vitro* **(Fig 8B)**. After 16 hours, TomatoRed(+) cells were isolated by FACS for analysis. To confirm expression of the key components of the WNT9b signaling pathway identified above, we extracted RNA from TomatoRed cells of 8 embryonic kidneys and analyzed transcripts levels (droplet digital PCR). As seen in Table 2, we confirmed mRNA expression of *Wt1, Fzd5, RSPO1* and *Lrp6* in the *Cited1*/Tomato(+) progenitor cells from E15.5 cap mesenchyme.

**Fig 8A:**
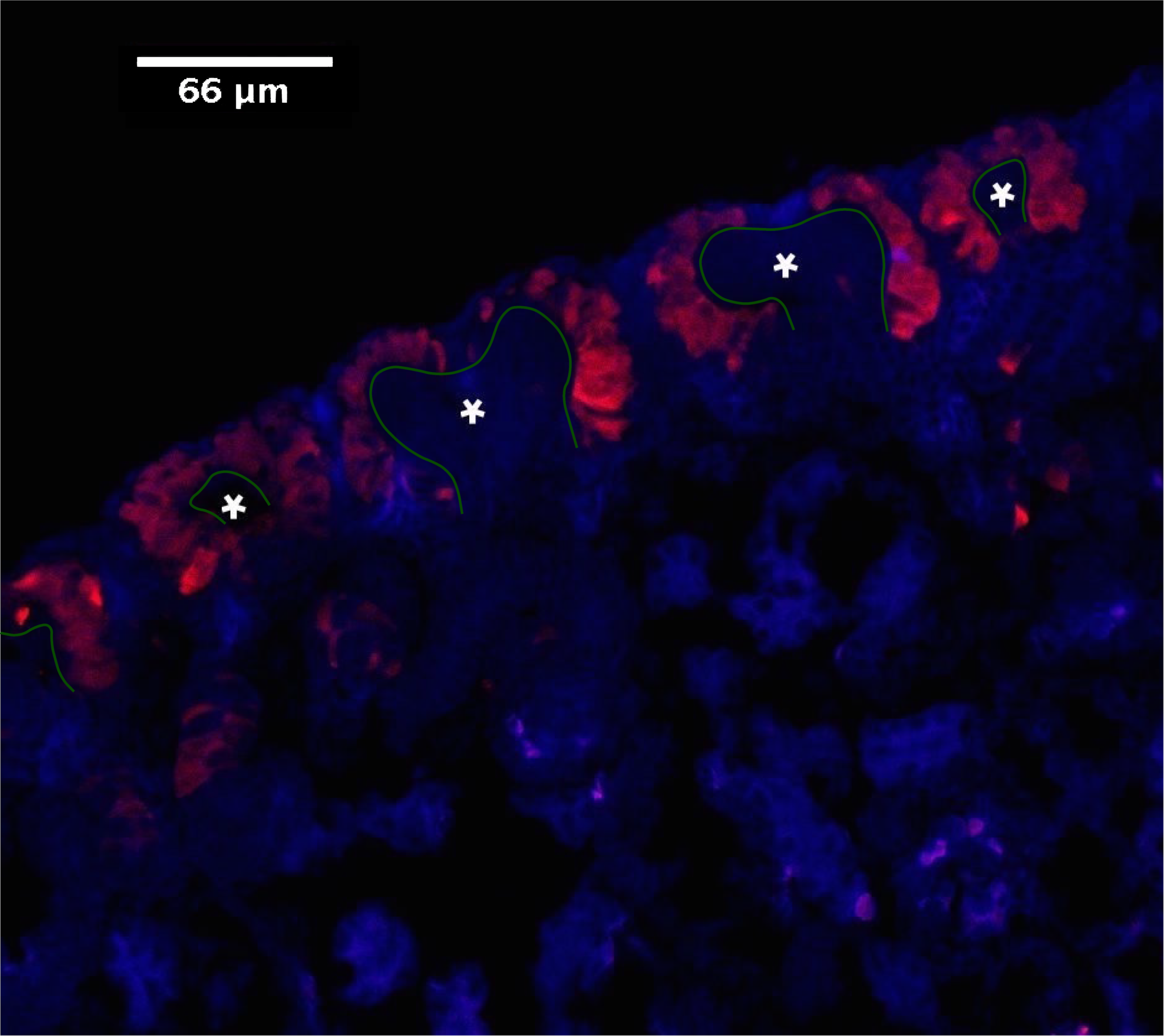
Identification and isolation of Citedl expressing cells from embryonic kidneys. A) Cryosections of E18 embryonic kidneys isolated from Citedl^Cre^/TomatoRed mice were assessed by immunofluorescent microscopy for the presence of TomatoRed in cap mesenchyme surrounding ureteric bud tips. (*) Ureteric Bud outlined in green. B) Whole E15.5 embryonic kidneys were dispersed into monolayer culture in the presence of tamoxifen (2.5 ng/ml) for 16 hours and TomatoRed(+) cells were visualized by immunofluorescent microscopy.

**Fig 8B:**
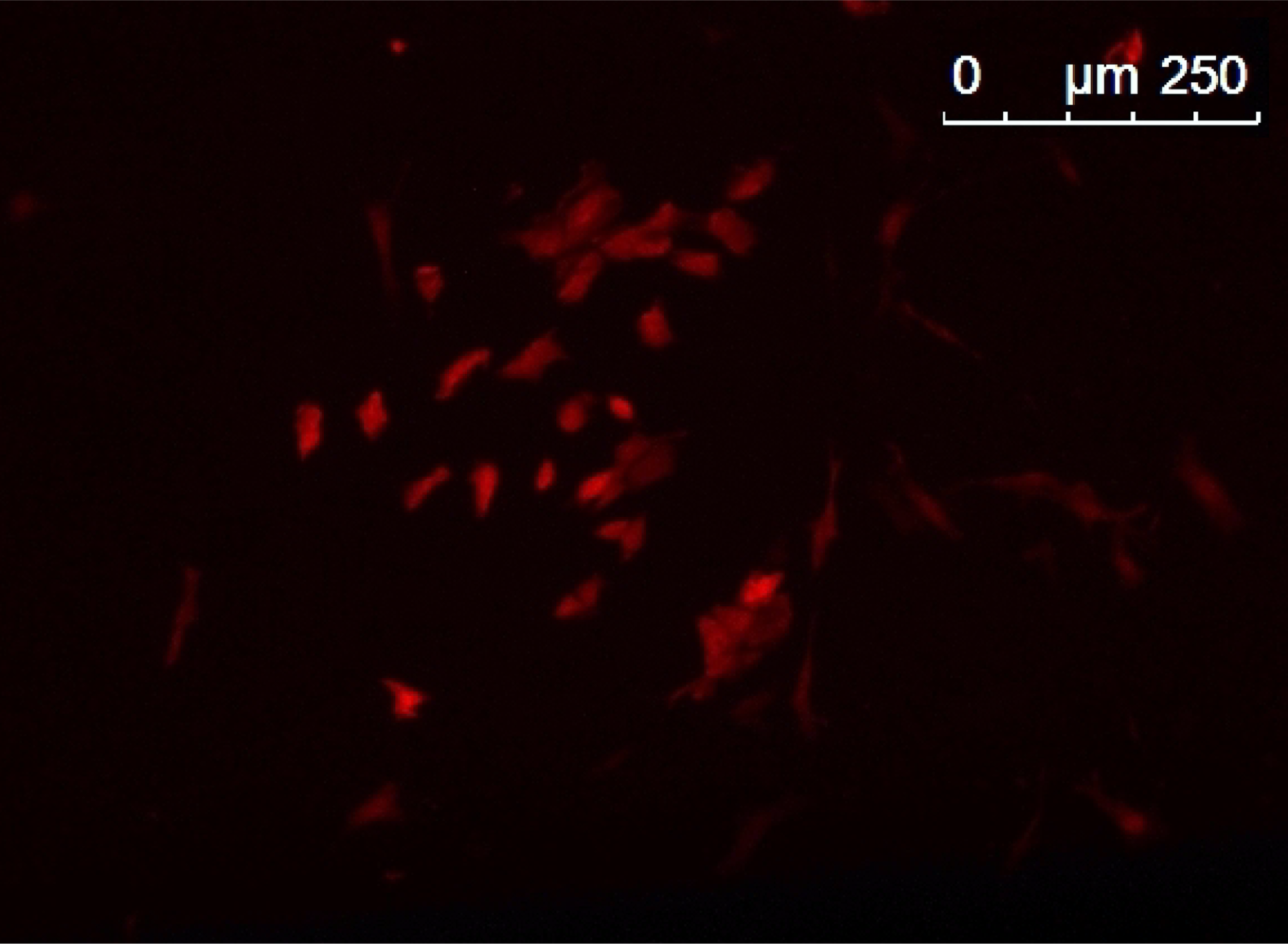
Identification and isolation of Citedl expressing cells from embryonic kidneys. Whole E15.5 embryonic kidneys were dispersed into monolayer culture in the presence of tamoxifen (2.5 /jg/ml) for 16 hours and TomatoRed(+) cells were visualized by immunofluorescent microscopy.

**Table II.**
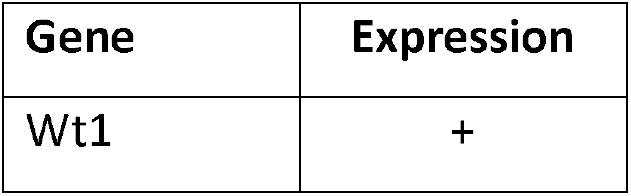
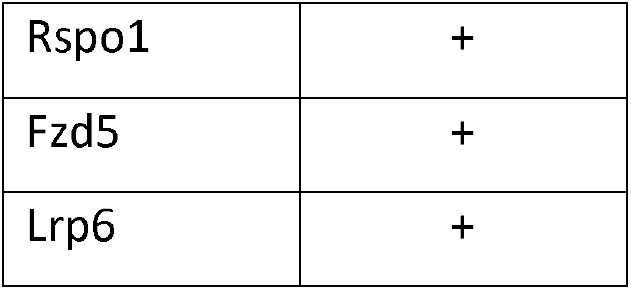
Expression of WNT9b pathway component transcripts in Citedl^Cre^/TomatoRed cells isolated by FACS from E15.5 embryonic mouse kidney.

| Gene | Expression |
| --- | --- |
| Wt1 | + |
| Rspo1 | + |
| Fzd5 | + |
| Lrp6 | + |

## Discussion

At about embryonic day E9.0 of mouse development, a lineage of WT1-expressing progenitor cells emerge within the OSR1(+) intermediate mesoderm. To model this early nephron progenitor cell (NPC) prior to the arrival of the ureteric bud, we studied the M15 cell line isolated from E10.5 mouse kidneys (Larsson et al., 1995). These cells express Osr1 mRNA and WT1 mRNA and protein, placing them in the NPC lineage. In previous studies from our lab, WT1 is essential for responsiveness to the inductive WNT9b signal, since it suppresses EZH2, a histone H3K27 tri-methylase, which in turn opens up chromatin permitting exit from the stem cell state (Aiden et al., 2010; Akpa et al., 2015). In this manner, WT1 was shown to activate β-catenin expression (Akpa et al., 2015). Surprisingly, however, we found that M15 cells were unresponsive to WNT9b in vitro. This suggests that WT1 expression alone is not sufficient to prime the NPC for WNT responsiveness and that the early NPC must acquire additional molecular properties by the time the ureteric bud arrives at E10.5-E11.

Although M15 cells are unresponsive to WNT9b, they express many components of the canonical WNT signaling pathway, including 4 frizzled receptors (Fzd1, Fzd2, Fzd3 and Fzd5) which can be demonstrated (in situ hybridization) in the cap mesenchyme surrounding each ureteric bud tip. M15 cells also express the frizzled co-receptor Lrp6 and complex-stabilizing proteins LGR4/6, shown by the GUDMAP consortium to be present in cap mesenchyme (McMahon et al., 2008; Harding et al., 2011). However, several investigators have shown that canonical WNT signal transduction is dramatically increased by stabilization of the Fzd/LRP6/WNT ligand at the cell surface (Binnerts et al., 2007). This requires a member of the R-spondin family which binds to the WNT-receptor complex through its association with an LGR family member (Carmon et al., 2011). The mechanism of R-spondin agonist activity was studied and shown to act through an interaction with ZNRF3. In complex with RNF43, ZNRF3 is a negative regulator of canonical Wnt-signalling and has a role of ubiquitinating Fzd receptors, targeting them for destruction and also preventing phosphorylation of Lrp receptors, keeping them in their inactive form (Koo et al., 2012; Hao et al., 2012). R-spondin1 transcripts are strongly expressed in the cap mesenchyme of E11.5 mouse kidney (Motamedi et al., 2014) but were entirely absent in M15 cells. In our study, pre-treatment of M15 cells with R-spondin1 enhanced WNT9b-induced canonical signaling activity fourfold. When in the cells were transfected with additional Fzd5, RSPO1 augmented the response to WNT9b elevenfold. Thus, RSPO1 appears to be critical for a robust response to WNT9b and its absence in M15 cells precludes measurable signal transduction. We postulate that RSPO1 is not expressed in the early E10.5 and the effects of WT1 on NPC chromatin alone are insufficient to induce RSPO1 expression. RSPO1 expression may be a late priming event in the maturation of the NPC.

The effects of RSPO1/LGR interactions are crucial for normal nephrogenesis. Three LGRs (4,5,6) interact with R-spondin proteins (Carmon et al., 2011; de Lau et al., 2011; Glinka et al., 2011; Gong et al., 2012; Ruffner et al., 2012). Lgr5 has been well-studied in intestinal epithelia where it is important for intestinal stem cell renewal (Barker et al 2007; Barker et al., 2010; Kinzel et al., 2014). In embryonic kidney, both Lgr4 and Lgr6 are expressed in the NPC lineage; we detected both Lgr4 and 6 transcripts in M15 cells. Since currently available siRNAs knockdown both Lgr4 and Lg6, we can’t determine which protein is most important in NPCs. However, fetal mice with knockout of the Lgr4 gene exhibit increased apoptosis of NPC and disturbance of the process by which NPC condense around ureteric bud tips (Mohri et al., 2012). This suggests that Lgr4 may be the primary determinant of WNT9b signal transduction in cap mesenchyme. In contrast, murine knockout of the *RSPO1* gene has no renal phenotype (Chassot et al., 2008), likely reflecting redundancy between RSPO1 and RSP03, both of which are expressed in the cap mesenchyme (GUDMAP). This is in keeping with our *in vitro* observations indicating that both recombinant RSPO1 and RSP03 enhance WNT responsiveness of M15 cells.

Few studies have investigated frizzled expression in the developing kidney. Ureteric bud specific expression of FZD4 and FZD8 in E11.5 kidneys was previously examined using Fzd4-lacZ and Fzd8-lacZ mouse models (Ye et al., 2011). Additionally, widespread renal expression of FZD2 and FZD7 was observed in 12, 13 and 18-week human fetal kidneys (Metsuyanim et al., 2009). Our in-situ hybridization data revealed distinct Frizzled expression patterns in E11.5 mouse kidneys. We observed *Fzd1, Fzd2, Fzd3, Fzd5* and *Fzd7* expression in the cap mesenchyme, whereas *Fzd4, Fzd6* and *Fzd8* expression was highly restricted to the ureteric bud. *Fzd10* expression was relatively non-specific and *Fzd9* in situ hybridization did not work for technical reasons. Interestingly, we found that in the presence of RSPO1, only *Fzd5* was limiting the WNT response. The canonical signal was amplified by transfecting cells with *Fzd5* but none of the other FZD family members. Furthermore, siRNA knockdown of *Fzd5* (but not *Fzd1* or *Fzd2)* reduced WNT9b responsiveness. These observations suggest that FZD5 is the primary WNT receptor involved in transducing the inductive WNT9b signal in mammalian kidney. Moreover, it raises the possibility that the other FZDs expressed in cap mesenchyme might be involved in transduction of other canonical and non-canonical WNT ligands, such as WNT6 and WNT11 from ureteric bud tips (Kispert et al., 1996; Itaranta et al., 2002) or WNT2b and WNT4 from the metanephric mesenchyme of the developing kidney (Stark et al., 1994; Lin et al., 2001).

Phylogenetic analysis of human frizzled proteins established five distinct frizzled subgroups (MacDonald and He, 2012), one of which consisted of FZD5 and FZD8. Our in situ hybridization studies of E11.5 embryonic mouse kidney demonstrate Fzd5 in cap mesenchyme, Fzd8 is exclusively expressed in the ureteric bud. Interestingly, WNT9b was demonstrated to bind and form a complex with FZD8 and Lrp6 (Bourhis et al., 2010). It is conceivable that Fzd8 mediates the robust canonical WNT signaling activity in ureteric buds reported by Bridgewater (Bridgewater 2008) and Iglesias (Bridgewater et al., 2008; Iglesias et al., 2014). Targeted excision of CTNNB1 in the ureteric bud blocked branching morphogenesis and bilateral renal aplasia (Bridgewater 2008).

Based on our data and the observations above, we propose a model of renal development in which WT1(+) NPC in E10.5 embryonic mouse kidney express some, but not all, components of the canonical WNT signaling pathway. By E11.5, additional events (expression of RSPO1 and increased expression of FZD5) have primed NPCs forming the cap mesenchyme, allowing responsiveness to the anticipated WNT9b signal from ureteric bud **(Fig 9).**

**Fig 9:**
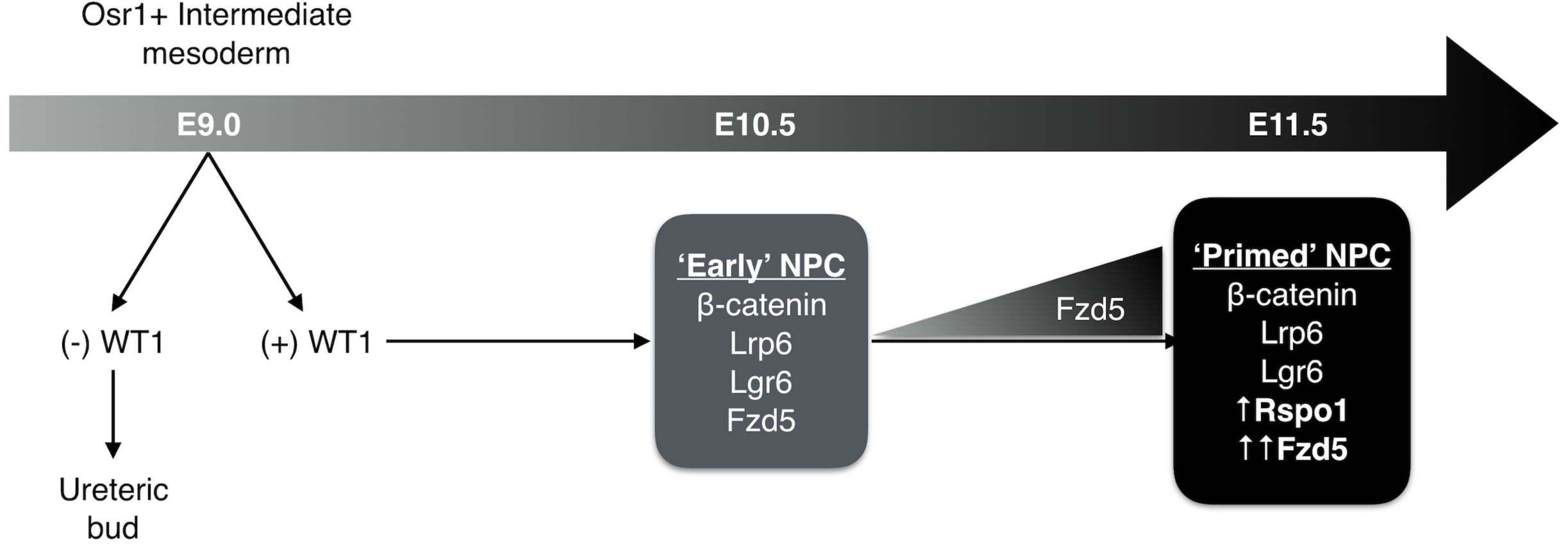
Proposed model of nephron progenitor cell development in embryonic mouse kidney Early WT1 (+) NPCs express a number of important molecules in the canonical WNT-signalling pathway. By E11.5, increased expression of Fzd5 and addition of RSPO1 render NPCs fully competent to respond to the inductive WNT9b signal.

## Methods

### Cell Culture

M15 cells (Larsson et al., 1995) are WT1-expressing cells isolated from E10.5 mouse mesonephric mesenchyme expressing the large T protein of polyoma virus under control of the early viral enhancer. Cells growing in monolayer attached to plastic culture vessels in the presence of DMEM culture medium with 10% Fetal Bovine Serum and 1% Penicillin/ Streptomycin.

### Luciferase Reporter transfections and Dual Luciferase Assay

Transient transfections were performed using a canonical WNT signalling reporter plasmid, Super8XTOPFlash. The Renilla luciferase expression vector pRL-SV40 (Promega, Madison, W1, USA) was used to normalize for transfection efficiency. Transfections for each condition were performed in triplicate and repeated three times on different days. The following frizzled plasmids were gifts from Chris Garcia & Jeremy Nathans: pRK5-mFzd1-1D4, pRK5-mFzd2-1D4, pRK5-mFzd3-1D4, pRK5-mFzd4-1D4, pRK5-mFzd5-1D4, pRK5-mFzd6-1D4, pRK5-mFzd7-1D4, pRK5-mFzd8-1D4, pRK5-mFzd9-1D4, pRK5-mFzd10-1D4 and pRK5-Wnt9b (Yu et al., 2012)(Addgene, Cambridge, MA, USA). Lrp5 (Clone ID: 3154246) and Lrp6 (Clone ID: 6409058) plasmids were purchased from Dharmachon (Lafayette, CO, USA).

One day prior to transfection, 20,000 M15 cells were seeded in 24-well plates and transfected at 80% confluency using Lipofectamine 2000 Transfection Reagent according to the manufacturer’s instructions (Thermo Fisher Scientific, Waltham, MA, USA). Plasmids were transfected in the following amounts: Fzd (50 ng), Super8XTOPFlash (44 ng), Lrp (5 ng), Wnt (50 ng), Renilla (1 ng). Recombinant Wnt9b (3669-WN/CF, R&D Systems, Minneapolis, MN, USA) was added at a concentration of 50 ng/mL to transfection media at the time of transfection in corresponding conditions. In R-spondin conditions, either 200 ng/mL of recombinant mouse RSPO1 (3474-RS - R&D Systems, Minneapolis, MN, USA) or 200 ng/mL of recombinant mouse Rspo3 (4120-RS/CF - R&D Systems, Minneapolis, MN, USA) was added to each well 24 hours post transfection. Firefly and renilla luciferase reporter activities were measured after 48h using the Dual Luciferase Assay System reagents and quantified in a GLOMAX 96 microplate luminometer (Promega, Madison, W1, USA). The reporter activity was expressed as a Firefly luciferase/ Renilla luciferase ratio.

The same procedure as described above was followed to monitor luciferase activity. For siRNA experiments, cells were transfected with Silencer^®^ pre-designed siRNA targeting mouse Fzd1 (siRNA ID: 75730), Fzd2 (siRNA ID: 57265), Fzd5 (siRNA ID: 14367) and Lrp6 (siRNA ID: 62715) (Ambion, Carlsbad, CA, USA) using Lipofectamine 2000 transfection reagent (Thermo Fisher Scientific, Waltham, MA, USA) according to manufacturer instructions.

### RNA isolation and Real-Time PCR Analysis

RNA was isolated using the QIAGEN RNeasy kit according to the manufacturer’s instructions (QIAGEN, Toronto, ON, Canada). RT-PCR was performed using the iScript cDNA synthesis kit (Bio-Rad, Mississauga, ON, Canada). Quantitative real-time PCR was performed using the SsoFast EvaGreen Supermix with Low ROX (Bio-Rad, Mississauga, ON, Canada) and specific primer sets in a LightCycler 480 II (Roche Applied Science, Laval, QC, Canada).

### Immunoblotting

Protein content was quantified in cellular extracts using the BCA assay (Pierce, Rockford, IL, USA). Twenty-five micrograms of protein extract were loaded onto SDS-PAGE gel and subjected to electrophoresis following standard immunoblotting techniques. The following primary antibodies and titres were used: anti-WT1 (antibody C19: sc-192, 1/200, Santa Cruz Biotechnology, Santa Cruz, CA, USA), anti-Actin (A5441, 1/10000, Sigma-Aldrich, Oakville, ON, Canada). Immunoreactive bands were detected using species-specific horseradish peroxidase-conjugated secondary antibodies (1/2000, Cell Signaling, Danvers, MA, USA) and visualized and analyzed using the GE Healthcare ECL Plus Western Blotting Detection Reagents and the BioRad Imager Scanner and software (GE Healthcare, Mississauga, ON, Canada).

### In situ hybridization

Section in situ hybridization of E11.5 embryos was performed according to the protocol listed on the GUDMAP website (https://www.gudmap.org/chaise/recordset/#2/Protocol:Protocol@sort(RID). cDNAs were purchased from ThermoFischer/Open Biosystems. For each gene, we include the clone ID, the restriction enzyme used to linearize the plasmid and the polymerase used to synthesize the antisense probe. Fzd1 (Clone ID: 5697795) Sall/T3, Fzd2 (Clone ID: 6411627) Sall/T3, Fzd3 (Clone ID: 30084926) EcoRI/T3, Fzd4 (Clone ID: 4238940) Sall/T7, Fzd5 (Clone ID: UI-M-CGOP-BRL-B-03-0-UI) EcoRI/T3, Fzd6 (Clone ID: 3983985) Sall/T7, Fzd7 (Clone ID: 6844727) Sall/T3, Fzd8 (Clone ID: 3992722) Sall/T7, Fzd9 (Clone ID: UI-M-CGOP-BGI-E-03-0-UI), Fzd10 (Clone ID: 556296) Pst1/T7.

### Mice

All animal experiments were approved by the McGill Facility Animal Care Committee (FACC). Cited1-Cre mice were donated from Dr. Mark de Caestecker (Boyle et al., 2008). Tom^flox/flox^ mice were bought from Jackson Laboratories. Cited1-Cre males were crossed with homozygous Tom^flox/flox^ females to generate double transgenic embryos. For immunofluorescence experiments, at 17 dpc, 0.1 mg/kg of Tamoxifen (Sigma) was administered to pregnant females via intraperitoneal injection. Females were sacrificed 24 hours later and embryos were harvested. For ddPCR experiments on Cited1/Tom cells, 2.5 μg/mL of 4-hydroxytamoxifen was administered to culture media in vitro after embryonic kidneys were digested in a collagenase B digestion solution at 37°C for 1 hour. Digested embryonic kidneys from one pregnancy were pooled and cells were grown at 37°C in tissue culture flasks in NPC growth media (Zhang et al., 2008).

### Tissue preparation and confocal microscopy

Embryonic mouse kidneys (E18) from Cited1/Tom mice were fixed overnight in *4%* PFA at 4°C. Kidneys were then transferred into 15% Sucrose in PBS and rocked at room temperature for 30 mins followed by rocking overnight at 4°C in 30% sucrose. Next, kidneys were placed into a 1:1 mixture of 30% sucrose/PBS and OCT and rocked at 4°C for 2 hours and then were embedded in OCT and stored at -80°C until sectioned. Cryosections (7uM) were obtained using a Leica Cryostat. Nuclei were counterstained with VECTASHIELD Antifade Mounting Medium with DAPI (Vector Laboratories, Burlingame, CA, USA). Images were obtained with a laser scanning confocal microscope (LSM780) and the ZEN2010 software (Carl Zeiss Canada Ltd., Toronto, ON, Canada) at room temperature and processed by Adobe Photoshop and Illustrator software.

### Fluorescence activated cell sorting (FACS)

Whole embryonic kidneys were isolated and activated with tamoxifen as previously described. Cells were then washed in PBS and resuspended into 500 μL of 2% FBS in PBS solution and kept at 4°C until they were sorted. Cell sorting was performed by immunophenotyping core facility staff using a BD FACSAria Fusion. Isolated Cited1/Tom cells isolated were immediately pelleted and frozen at -80°C.

### Droplet digital PCR (ddPCR)

RNA was extracted from Cited1/Tom cells followed by cDNA synthesis as previously described (n=4). Droplets were formed in a QX200 Droplet Generator and PCR was performed using the QX200 ddPCR EvaGreen Supermix (Bio-Rad, Mississauga, ON, Canada) and specific primer sets in a C1000 Touch Thermal Cycler (Bio-Rad, Mississauga, ON, Canada). Droplets were read using the QX200 Droplet Reader machine and results were displayed in QuantaSoft software.

### Statistical Analysis

Graphs are presented as mean ± s.e.m of three or more independent results. Statistical significance was assessed by a one-way ANOVA followed by a Dunnett correction for multiple comparisons.

## Acknowledgements

The authors would like to acknowledge background information drawn from the GUDMAP consortium public database (www.GUDMAP.org). We would like to acknowledge coinvestigators of the CIHR/FRSQ/ERARE consortium who gave advice and critical analysis of the manuscript.

## Competing Interests

No competing interests declared

## Funding

This work was supported by operating grants to Dr. Goodyer from the Kidney Foundation of Canada, the Canadian Institutes of Health/ Fonds de recherche du Québec - Santé/ERA-Net for Research Programs on Rare Diseases and an infrastructure support grant to the McGill University Health Center Research Institute from the Fonds de la Recherche en Santé du Québec (FRSQ). Kyle Dickinson was the recipient of a graduate studentship award from the Research Institute of McGill University Health Centre and Desjardins Group. Paul Goodyer is the recipient of a CIHR James McGill Research Chair.

## Data Availability

Not Applicable

## FIGURE LEGENDS

**Fig 1. WT1 is expressed in MIS cells. A)** mRNA from E10.5 mouse mesonephric mesenchyme (M15 cells) was analyzed by RT-PCR for *Wt1* mRNA expression in M15 cells and MK3 (positive control) (Potter) cells vs water blank. B) Lysates of M15 cells vs E14.5 MK4 (negative control) (Potter) or am318.2 mesenchymal stem cells from 20-week gestation human amniotic fluid (Murielle) were analyzed by Western immunoblotting for WT1 protein (upper panel) and Beta actin (lower panel).

**Fig 2. WNT-responsiveness of M15 cells.** M15 cells were transiently transfected with **β-** catenin-luciferase reporter (8X SuperTOPFlash) and Renilla-luciferase reporter. The cells were exposed to recombinant WNT9b (50 ng/ml). After 48 hours, TOPFlash to Renilla signal (RLU) was measured in a luminometer. An unpaired two-tailed Welch’s t-test was performed, (ns) p=0.98.

**Fig 3. Effect of recombinant RSPO1 on responsiveness of** M15 **cells to WNT9b.** M15 cells were transfected with 8X SuperTOPFlash, Renilla and *Wnt9b* plasmids and cultured for 24 hours; recombinant RSPO1 or RSP03 (200 ng/ml) were added for an additional 24 hours and TOPFlash to Renilla signal was measured. A one-way ANOVA followed by a Dunnett correction for multiple comparisons was performed. (****) = p <0.0001

**Fig 4. Frizzled (FZD) mRNA expression in embryonic day E11.5 mouse kidney.** Cryosections of E11.5 mouse kidneys were assessed by *in situ* hybridization using riboprobes for *Fzd 1-10,* except *Fzd9* which was unsuccessful for technical reasons.

**Fig 5. Effect of FZD expression on WNT9b responsiveness in** M15 **cells.** A) M15 cells were transiently transfected with β-catenin-luciferase reporter (8X SuperTOPFlash), Renilla-luciferase reporter, l/1/nt9jb-expression vector and various *Fzd* 1-10 expression plasmids in the presence of recombinant RSPO1 (200 ng/ml). TOPFlash to Renilla signal was measured after 48 hours. A one-way ANOVA followed by a Dunnett correction for multiple comparisons was performed. (**) p=0.0002 B) M15 cells were transiently transfected with (3-catenin-luciferase reporter (8X SuperTOPFlash), Renilla-luciferase reporter, *Wnt9b-express\on* vector and siRNAs targeting *Fzd*1, *Fzd2* or *Fzd5* vs a scrambled negative control siRNA in the presence of recombinant RSPO1 (200 ng/ml). TOPFlash to Renilla signal was measured. A one-way ANOVA followed by a Dunnett correction for multiple comparisons was performed. (*) p=0.005

**Fig 6. LRP6 is required for optimal responsiveness of M15 cells to WNT9b**. M15 cells were transiently transfected with (3-catenin-luciferase reporter (8X SuperTOPFlash), Renilla-luciferase reporter and a *Wnt9b* expression vector and treated with RSPO1 (200 ng/ml). The cells were cotransfected with an siRNA targeting *Lrp6* or a scrambled negative control siRNA in the presence of recombinant RSPO1 (200 ng/ml). After 48 hours, TOPFlash to Renilla signal was measured. In another experiment, the cells were co-transfected with an *Lrp5* expression plasmid to assess its effect on WNT9b pathway activity. A one-way ANOVA followed by a Dunnett correction for multiple comparisons was performed. (****) pcO.OOOl

**Fig 7. Responsiveness to extrinsic WNT9b is restored by addition of *Fzd* and RSPO1.** In all conditions, M15 cells were transiently transfected with β-catenin-luciferase reporter (8X SuperTOPFlash) and Renilla-luciferase reporter; in some experiments the cells were cotransfected with *FzdS* expression plasmid. TOPFlash to Renilla signal was measured in the presence or absence of recombinant WNT9b (50 ng/mL) and/or recombinant RSPO1 (200 ng/ml). A one-way ANOVA followed by a Dunnett correction for multiple comparisons was performed. (****) = pO.OOOl

**Fig 8. Identification and isolation of Cited1 expressing cells from embryonic kidneys.** A) Cryosections of E18 embryonic kidneys isolated from Cited1^Cre^/TomatoRed mice were assessed by immunofluorescent microscopy for the presence of TomatoRed in cap mesenchyme surrounding ureteric bud tips. (*) Ureteric Bud outlined in green. B) Whole E15.5 embryonic kidneys were dispersed into monolayer culture in the presence of tamoxifen (2.5 ng/ml) for 16 hours and TomatoRed(+) cells were visualized by immunofluorescent microscopy.

**Fig 9. Proposed model of nephron progenitor cell development in embryonic mouse kidney.** Early WT1(+) NPCs express a number of important molecules in the canonical WNT-signalling pathway. By E11.5, increased expression of Fzd5 and addition of RSPO1 render NPCs fully competent to respond to the inductive WNT9b signal.

